# Somatic Evolution Introduces Distinctive Changes in Proteome Composition in Cancer Types

**DOI:** 10.1101/451310

**Authors:** Viktoriia Tsuber

## Abstract

On the proteome level, somatic evolution in cancer is asymmetric, with several amino acids being irreversibly lost and the others gained across cancer types. However, though the gains and losses of amino acids are consistent in multiple cancer types, their magnitudes and prevalences differ. Here, we analyzed data on circa three million amino acid substitutions in the Catalogue of Somatic Mutations in Cancer (COSMIC) database to investigate profiles of gains and losses of amino acids in 23 human cancer types. We found characteristic differences in amino acid gain/loss profiles in the cancer types, with several of them demonstrating very specific profiles of proteome alterations associated with distinct endogenous or exogenous factors.

## Introduction

Cancer arises in somatic evolution in tissues when a cell acquires a sufficient number of “driver” mutations that promote its unrestrained proliferation, growth, dissemination and invasion. Yet, most mutations in the cancer cell are “passenger” mutations that accumulate during the cell’s lifetime in a buildup of DNA damage. Latest advances in comprehensive sequencing, the genome-wide association study approach and the worldwide effort in creation of large public cancer databases have granted significant progress in studies of mutational landscape in cancer. Recent profound analyses of nucleotide alteration signatures attributed mutational profiles to effects of external and internal deleterious factors in different cancer types.^1, 2^

Nonsynonymous substitutions in the nucleotide sequence of a protein-coding part of a gene ultimately result in changes of the amino acid composition of the protein product of the gene, and consequently in changes of the structure and properties of the protein. However, though mutations in individual cancer driver proteins are broadly studied, few studies only have examined the global landscape of amino acid substitutions in cancer.^3,4^ We have recently shown, that somatic evolution in cancer consistently changes composition of the cancer cell proteome, where some amino acids permanently disappear from, and others are *de novo* introduced into proteins across multiple cancer types.^4^ We found, that Arg is by far the most prominent irreversible loss and His and Cys are the most substantial *de novo* gains in proteins in cancer, but other amino acids are also steadily lost or gained, thus making somatic evolution in cancer asymmetric. In the total cancer proteome, Ala, Asp, Glu, Gly, Pro and Ser universally disappear from the protein composition, while Phe, Ile, Lys, Leu, Met, Asn, Gln, Thr, Val, Trp, Tyr are almost constantly gained. However, the magnitudes of the amino acid gains/losses vary extensively between cancer types, thus creating distinctive molecular profiles of their somatic evolution.

In the present study, we analyzed profiles of mutation-induced alterations of the proteome composition in 23 human cancer types, using data on over three million amino acid substitutions in twenty-one thousand cancer samples from the Catalogue of Somatic Mutations in Cancer (COSMIC; v81) database.^5^ We show that in a majority of the cancer types, their tissue-specific somatic evolution introduces characteristic changes into composition of proteins. We found that within-cancer similarities of amino acid gains and losses in samples vary between the cancer types, with the most homogeneous profiles in skin, cervical and endometrial cancers, and most heterogeneous profiles in kidney, ovarian and bone cancers. We demonstrate that the cancer-specific amino acid gain/loss signatures can be recognized with a machine learning algorithm in several cancer types.

## Materials and Methods

We extracted information on nucleotide substitutions and resulting amino acid substitutions from the COSMIC (v81) database (http://cancer.sanger.ac.uk). The analysis of the data was performed in R version 3.3.1^6^ using RStudio version 0.99.893.^7^

## Results

From the COSMIC v81 database,^5^ we extracted data on amino acid substitutions in proteins that arise from single nucleotide substitutions in the total of twenty-one thousand cancer samples. Missense and nonsense single nucleotide substitutions make a majority of the data in the COSMIC database, comprising about 93% of entries. Of these, nonsynonymous substitutions are 75%. We analyzed data on over three million amino acid substitutions for 23 cancer types, for which the number of tumor samples exceeds one hundred in the COSMIC database. The number of tumor samples ranged between 180 (cervical cancer) and 2807 (hematopoietic and lymphoid tissue cancers). The median number of the mutations in each tissue ranged from 9 (adrenal gland) to 216 (skin). We subtracted the number of lost amino acid residues from the number of introduced residues and thus obtained net numbers of gains/losses of amino acids. In this way, we calculated net gains/losses of amino acids for the cancer types in the analysis, all samples, and in some cases individual proteins in the samples. With the obtained net amino acid gain/loss data, we performed correlation analyses and hierarchical clustering analyses within individual cancer types. Besides, as different cancer types inherently have different mutation rates, we used percentages of the net gain/loss numbers of amino acids to the total number of the mutations in the cancer type for between-cancer type comparisons.

In the total cancer proteome, consistent patterns of net gains/losses of all amino acids are observed (Figure 1, Figures S1-S5). While the changes are virtually unidirectional across the cancer types, with few exceptions, the ranges of the gains/losses are different in individual cancer types (Figures S6-S11). A number of cancer types demonstrate extremely high gains or losses of several amino acids thus suggesting presence of very specific profiles of proteome alterations. For instance, skin cancer shows exraordinarily high gains of Phe, Lys, Leu and Gln, while losing greatly more Pro or Gly than any other cancer type. Similarly, lung cancer acquires elevated quantities of Leu and predominantly loses Gly, though to a smaller extent than skin cancer. Cancers of cervix and urinary tract demonstrate high losses of Asp, Glu and Ser and simultaneous elevated gains of Lys. Interestingly, while most cancer types gain Phe, endometrial cancer loses it. Furthermore, endometrial cancer gains Ile and Tyr at a much higher rate than other cancers do. Notably, skin cancer is the only cancer type, where His, Met and Trp are lost (Figure 2). In addition, upper aerodigestive tract cancers gain Gly and Pro, while all other cancer types lose the amino acids. Liver cancer gains higher percentage of Val compared to other cancer types. Figure 3 summarizes prevalences of rates of gains and losses of amino acids in each of the 23 cancer types.

**Figure 1.**
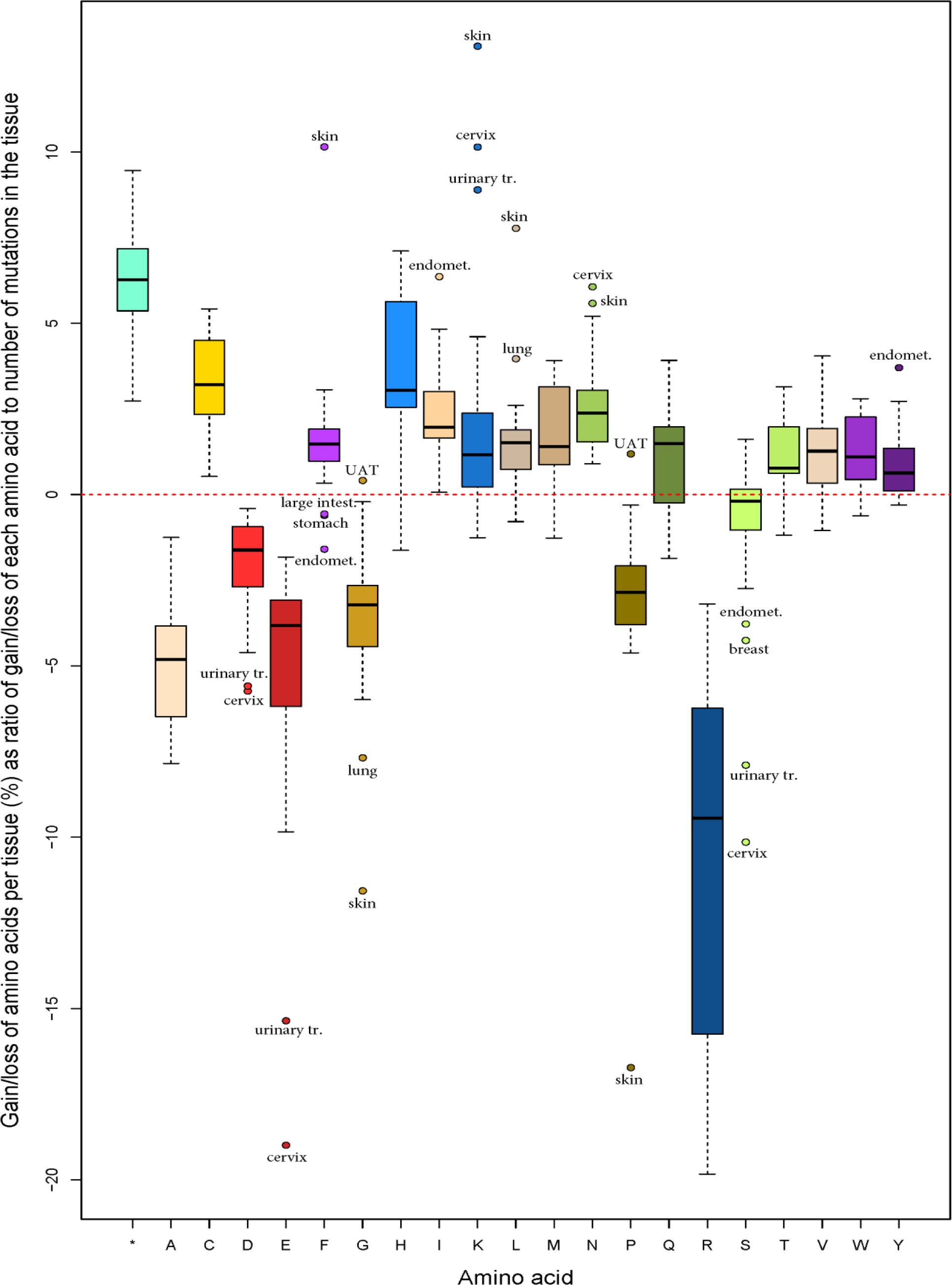
Mutation-induced changes of amino acid quantities in proteins in 23 major human cancer types as percentages of gains/losses of amino acids to the total number of the mutations in each cancer type. In this and the following graphs, amino acids are denoted with a one-letter code, and truncation is denoted as “*”; “UAT” denotes upper aerodigestive tract tumors.

**Figure 2.**
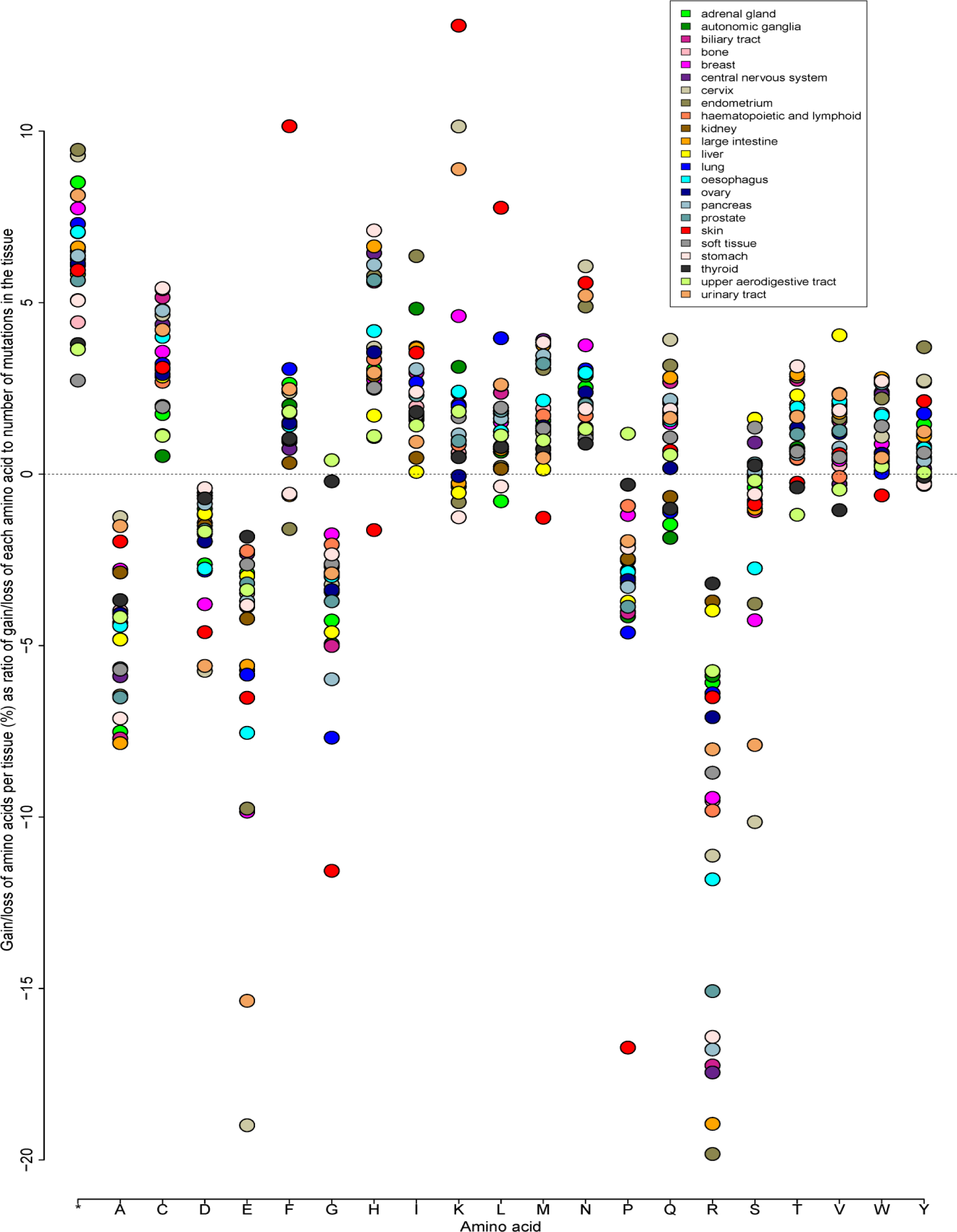
Rates of gains/losses of amino acids in proteins in the 23 cancer types.

**Figure 3.**
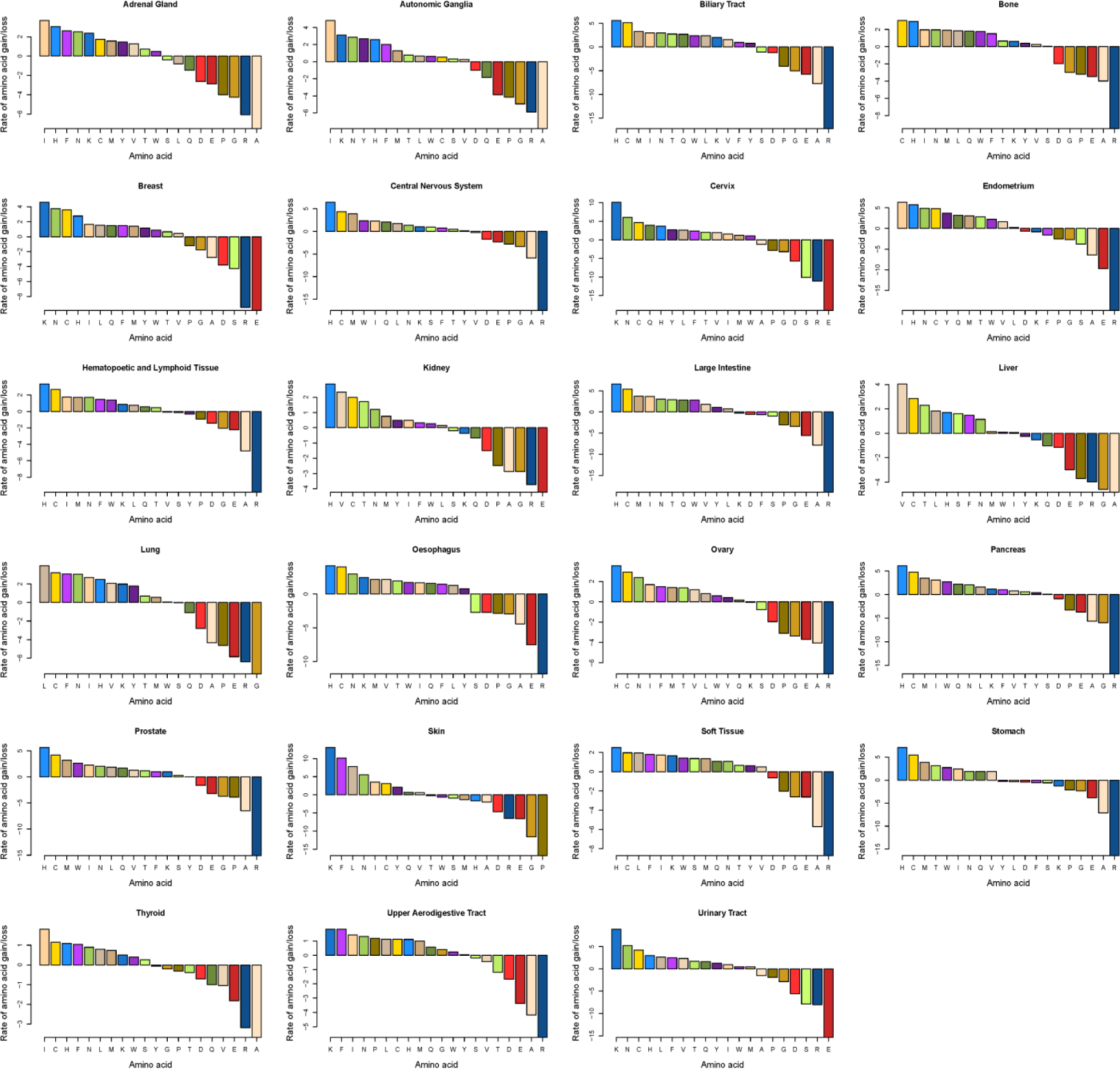
Rates of gains/losses of amino acids in proteins in each cancer type.

### Case studies of skin, cervical and endometrial cancers

We analyzed gains/losses of amino acids on the level of individual proteins in cancers of skin, endometrium and cervix. For each cancer type, we calculated numbers of proteins that undergo net gains/losses of its five most frequently gained and most frequently lost amino acids (Figure 4). For instance, the protein coded by the *UBR4* gene gains Phe residues thirty times more often than it loses them in all skin cancer samples combined, and is therefore counted as having gained Phe. In this way, we evaluate the scope of the proteome alterations in each of the three cancer types. In individual tumor samples, the numbers of proteins that undergo net changes of their amino acid composition depend on the severity of the tumor mutation load.

**Figure 4.**
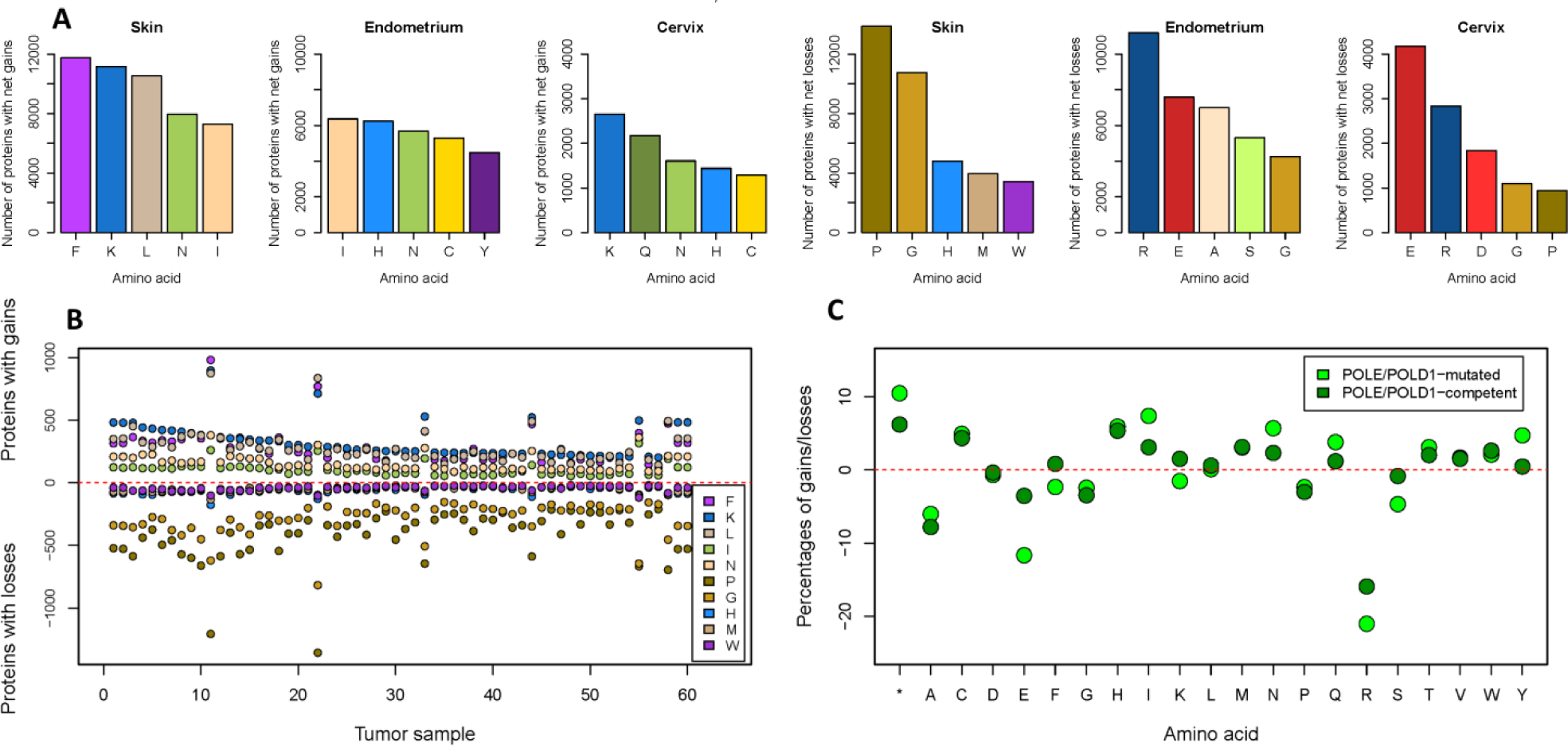
**A.** Numbers of proteins with gains of five most gained amino acids and with losses of five most lost amino acids in cancers of skin, cervix and endometrium. **B.** Numbers of proteins with gains of five most gained amino acids and with losses of five most lost amino acids in most mutated samples of skin cancer. **C.** Gains/losses of amino acids in *POLE/POLD1*-mutated and *POLE/POLD1*-competent tumors in endometrial cancer as percentages of the gains/losses of amino acids to the total number of mutations in the cancer subtypes.

#### A. Skin cancer

Of 966 samples of skin tumors in the database, 87.2% is malignant melanoma. For skin tumors, the percentage of the AT pair transformations in coding mutations is much smaller than that in samples from all cancer types combined, 9.8% in skin cancers vs 22.6% in the total COSMIC database, and second smallest of the 23 cancer types. It is consistent with prevalence of C>T mutations in skin cancers caused by ultraviolet-induced formation of pyrimidine dimers in DNA. In the proteome of skin cancer, gains of Phe, Lys, Leu, Ile and Asn and losses of Gly, Pro, His, Met and Trp are observed in a substantial portion of cell proteins (Figure 4A). For instance, Phe is *de novo* introduced in fifteen times more individual proteins than it is lost (Table S1) thus indicating the high tendency of skin cancer proteins to avidly gain the amino acid. Similarly, the number of proteins that irreversibly lose Gly is twelve times bigger than the number of those that *de novo* gain it. In skin cancer, proteins most affected by the gains and losses were predominantly those heavily mutated in many cancer types, such as the proteins coded by the genes *TTN*, *MUC16*, *PCLO*, *CSMD* etc. (Table S1). The proteins are often extremely large or coded for by late replicating genes.^2^ Several known cancer suppressor and oncogenic proteins demonstrate high numbers of gains/losses of some amino acids but not of others. For instance, the protein coded by the *TP53* gene comes in top ten of the proteins that lose Pro and His, but not Gly, Met or Trp. His is also actively lost in the group of zinc finger proteins that account for 6.3% of all proteins losing His vs 2.8% of proteins losing Pro. Interestingly, the p53 protein gains Trp against the prevalent tendency of losing the amino acid in skin cancer (Table S1). In highly mutated skin tumors, the gains and losses can potentially affect considerable numbers of individual proteins (Figure 4B). Arguably, such tumors cannot forgo synthesis of all the affected proteins and may have an increased demand for constant supply of some amino acids compared to normal tissue.

The predominant gains of the three hydrophobic acids Phe, Leu and Ile by the proteome of skin cancer may increase its hydrophobicity, as a concomitant loss of hydrophobic Trp and Met is much smaller. Furthermore, the proteome of skin cancer may become more basic due to a substantial gain of Lys. While His also has a basic side chain, its pKa is much smaller, being 6.04 vs 10.54 in Lys. Moreover, the loss of His is smaller than the gain of Lys in skin cancer.

#### B. Endometrial cancer

Of 273 endometrial cancer samples in the COSMIC database, 90.8% of tumors is defined as endometroid carcinoma, and 8.4% as serous carcinoma. There were two distinct subsets of tumors in the data on endometrium cancer. 37 samples contained mutations in *POLE* and *POLD1* genes coding for DNA polymerases Pol δ and Pol ∊ that are essential for faithful replication of DNA. The mutations are known to to cause proofreading errors and thus hypermutator phenotype.^8^ Indeed, the median number of mutations in the *POLE/POLD1-* mutated samples was 1515, and the data on the samples comprised 76.3% of the analyzed data for endometrial cancer. The median number of mutations in tumors with intact *POLE* and *POLD1* genes was 84. The two types demonstrated differences in their amino acid gain/loss profiles. The five most gained amino acids in *POLE/POLD1*-mutated tumors were Ile, His, Asn, Cys and Tyr. In *POLE/POLD1*-competent tumors, the five most gained amino acids were His, Cys, Met, Ile and Trp, while there was almost no gain of Tyr (Figure 4C). More truncation events and much bigger loss of Arg are also observed in *POLE/POLD1*-mutated tumors than in any other cancer type in the study, with more than 20% of coding mutations in samples of the cancer type being losses of Arg (Figure 4C). Gains of the five most gained amino acids Ile, His, Asn, Cys and Tyr affect similar numbers of proteins, while the number of proteins losing Arg is considerably higher than of those losing Glu, Ala, Ser and Gly in *POLE/POLD1*-mutated tumors (Figure 4A). Both *POLE/POLD1*-mutated and *POLE/POLD1*-competent tumors demonstrate high losses of Arg and substantial gains of Cys and His.

The alterations are consistent with the most prevalent substitutions of Arg being those by Cys, His, Gln, Trp and truncations.^3,4^ The exceptionally high irreversible loss of Arg is associated with the codons for the amino acid that contain the CG sequence in their composition.^4^ The codons are most likely affected by spontaneous deamination of methylated cytosines in CpG sequences^9^ that is observed in many cancer types of epithelial origin with high cell turnover.^10^ All samples of endometrial cancer were used to calculate numbers of proteins that undergo gains/losses of its most frequently gained and frequently lost amino acids. The tumor suppressor proteins p53 (28.9% tumor samples contain at least one mutation in the gene coding for the protein) and FBXW7 (17.6% samples) demonstrate extraordinarily frequent losses of Arg with simultaneous gains of Cys residues. Loss of Arg is an important alteration in the p53 protein where the amino acid residues participate in interactions with DNA, and it thus has an exceptionally high ratio of events of irreversible loss of Arg to the number of Arg residues in its molecule in cancer mutations.^4^ However, in skin cancer (16% samples) p53 is less affected by the loss of Arg, than in endometrial cancer. Loss of Arg constitutes 29% of mutations in p53 in skin cancer, vs 38.4% of that in endometrial cancer (chi-squared = 391.7; *p* < 2.2 × 10^−16^; Cramer’s V = 0.31). The finding may indicate that the amino acid gain/loss profile in mutations may have an effect on emergence of a cancer-driving mutated gene. It has been shown that mutation spectra of driver genes in cancer show high similarity to the tissue-specific adult stem cell mutation spectra.^11^ The PIK3CA protein (49.1% samples) most frequently loses Glu in positions 542 and 545, and the KRAS protein (20.9% samples) demonstrates high losses of Gly because of the gain-of-function mutation where Gly residues in positions 12 and 13 are replaced with another amino acid, comprising 0.4% and 0.9% of all losses of Glu and Gly, respectively. Interestingly, despite the tendency of proteins in endometrial cancer to lose Ala irreversibly, the giant protein titin coded for by the gene *TTN*, does not lose the amino acid at all. Titin is mutated in 107 samples in the endometrial cancer data. The protein tops the list of proteins losing Arg, Glu, Ser, and only comes second to the KRAS protein in losing Gly, but does not show loss of Ala. The amino acid composition of titin contains similar numbers of Gly and Ala, and the numbers of proteins that lose them are comparable within the cancer type.

#### C. Cervical cancer

178 of 180 samples of cervical cancer in the COSMIC database are defined as squamous cell carcinoma. Cervical cancer shows a remarkable preference for transformations of the GC nucleotide pair. The percentage of the AT nucleotide pair transformations is 8.1% that is the lowest value among all the cancer types. Cervical cancer is caused in almost all cases by human papillomavirus,^12^ and frequent mutations that change C to T or C to G are associated with activity of APOBEC family of cytidine deaminases that are innate immunity enzymes attacking retroviruses.^13^ Glu, Arg, Asp, Gly and Pro are five amino acid residues that are most frequently irreversibly lost in somatic evolution of cervical cancer. Lys, Asn, Gln, His and Cys are five most frequently gained amino acid residues. Similarly to endometrial cancer, the tumor suppressor protein FBXW7 (10% of samples are affected by at least one amino acid substitution in the protein) loses Arg and gains Cys, the PIK3CA protein (28.3% samples) loses Glu and gains Lys, and the KRAS protein (5.6% samples) loses Gly.

### Correlation analyses and hierarchical clustering analyses within 23 cancer types

We performed correlation analyses and hierarchical clustering analyses of net amino acid gains/losses for each tissue. Supposedly, Pearson correlation coefficients *r* that are close to |1| indicate that respective amino acids are constantly gained/lost together or either one of the amino acids is lost and the other is gained at equal rates in most samples in the cancer type. Consistently high correlation coefficients of gains and losses of amino acids in a cancer type indicate presence of a uniform amino acid gain/loss signature caused by a same mutational process in majority of tumor samples, while low values of *r* show heterogeneity of tumor samples and presence of a wider variety of underlying molecular processes. We calculated sums of positive and negative correlation coefficients in correlation matrices for each cancer type. The obtained amino acid gain/loss correlation scores may serve as indices of consistency of a molecular signature throughout a cancer type (Figure 5). Figure 6 shows correlation matrices for cancers of skin and kidney that have highest and lowest correlation scores, respectively. For skin tumors, a majority of *r* values is either bigger than 0.9 or smaller than −0.9. For instance, a change in number of Phe residues is strongly positively correlated to that of Lys, Leu and the total number of mutations (*r*=0.98), and strongly negatively correlated to that of Pro (*r*=−0.99). Interestingly, Pro is not directly substituted by Phe but continually substituted with Ser and Leu. In turn, Phe most frequently arises in place of Ser. The only weak association is that of a change in number of Thr residues with those of all other amino acids. Consistently, the gained and lost amino acids in skin cancer are grouped in very distinct clusters, with minimal distances between amino acids within each cluster and Thr solely separated from other amino acids (Figure 5). To the contrary, the amino acid gain/loss correlation matrix of kidney cancer demonstrates absence of strong associations, with the highest *r* value of 0.72 for the correlation between the total number of mutations and number of truncation events. Concurrently, big distances between amino acids in the cluster analysis diagram show inconsistencies of their associations. Though 60% of kidney tumor samples in the COSMIC database are classified as clear cell carcinomas, the operating mutational processes are probably very diverse and lead to different amino acid gain/loss outcomes in each case. The cancer types with next highest amino acid gain/loss correlation scores are those of endometrium, cervix, liver and large intestine (Table S2 summarizes correlation matrices for all the cancer types), while the cancer types with lowest correlation scores are those of autonomic ganglia, ovary, bone and central nervous system.

**Figure 5.**
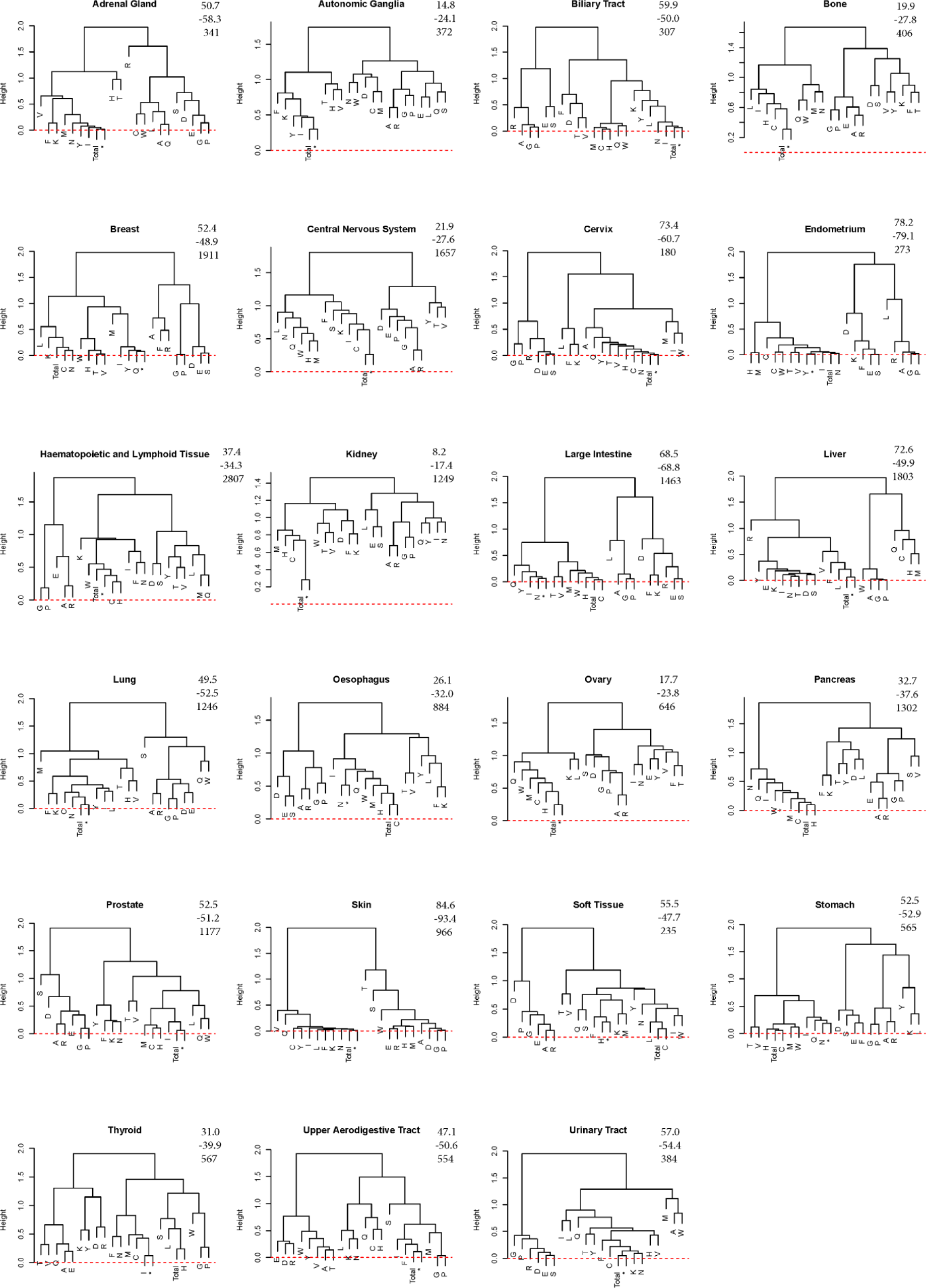
Hierarchical clustering diagrams of amino acid gains/losses in 23 cancer types. The numbers in the right upper corner of each diagram denote the positive correlation score, negative correlation score and number of samples of the cancer type in the COSMIC v81 database, respectively.

**Figure 6.**
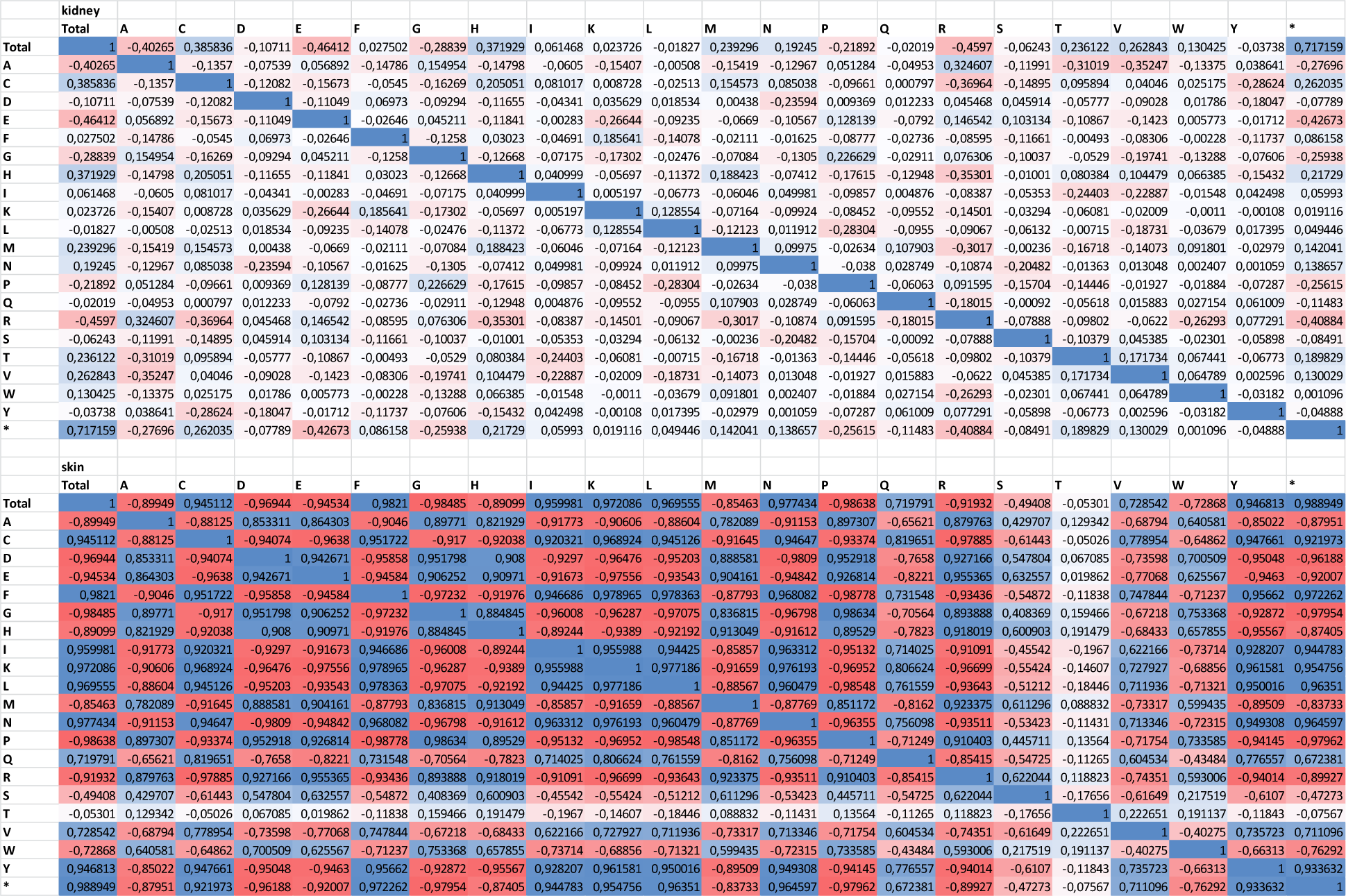
Amino acid gain/loss correlation matrices for cancers of kidney and skin. “Total” denotes the total number of mutations in the tissue.

### Machine learning analysis

We divided data on each cancer type into quartiles based on the number of mutations in tumor samples. For the quartiles of most heavily mutated samples in each cancer type, we performed a machine learning (ML) analysis of gains/losses of amino acids using the K-nearest neighbor method. We found that the ML analysis could indeed distinguish among different cancer types from their amino acid gain/loss profiles, albeit with unequal sensitivity. Skin, large intestine and lung cancers were the most accurately classified tissue types (Figure 7), with as much as 95% cases of skin cancer being correctly attributed. We suggest that the accuracy of the classification depends on the prominence and homogeneity of the cancer type-specific signature, number of analyzed tissue samples, and median number of mutations in the cancer type. For instance, skin cancer has a very distinct signature, uniformity of amino acid gains/losses throughout the samples, the highest median number of mutations of all cancer types, and a substantial though not the biggest number of samples in the database. Alhough amino acid gain/loss signatures of lung and large intestine cancers seem to be less divergent from those in other tissues than that of skin cancer (Figure 2), the cancer types are detected with 83% and 64% accuracy, respectively. The both cancer types are represented with high numbers of samples in the database, mostly have many mutations per sample and high amino acid gain/loss correlation scores (Figure 5). However, cancers of the central nervous system and those of hematopoietic and lymphoid tissue are reasonably well detected with the ML (56% and 58% accuracy, respectively) despite low median numbers of mutations and low to medium correlation scores. Interestingly, only 23% cases of breast cancer were correctly attributed despite its having the second biggest number of cases in the database and moderately high amino acid gain/loss correlation score. Accuracy of detection of breast cancer is thus smaller than that of cervical cancer with a tenfold smaller number of samples. The finding may indicate heterogeneity of breast cancer types due to a bigger variety of underlying molecular processes compared to those of skin or lung cancers. Remarkably, the ML model failed to detect soft tissue, biliary tract, bone and ovarian cancers.

**Figure 7.**
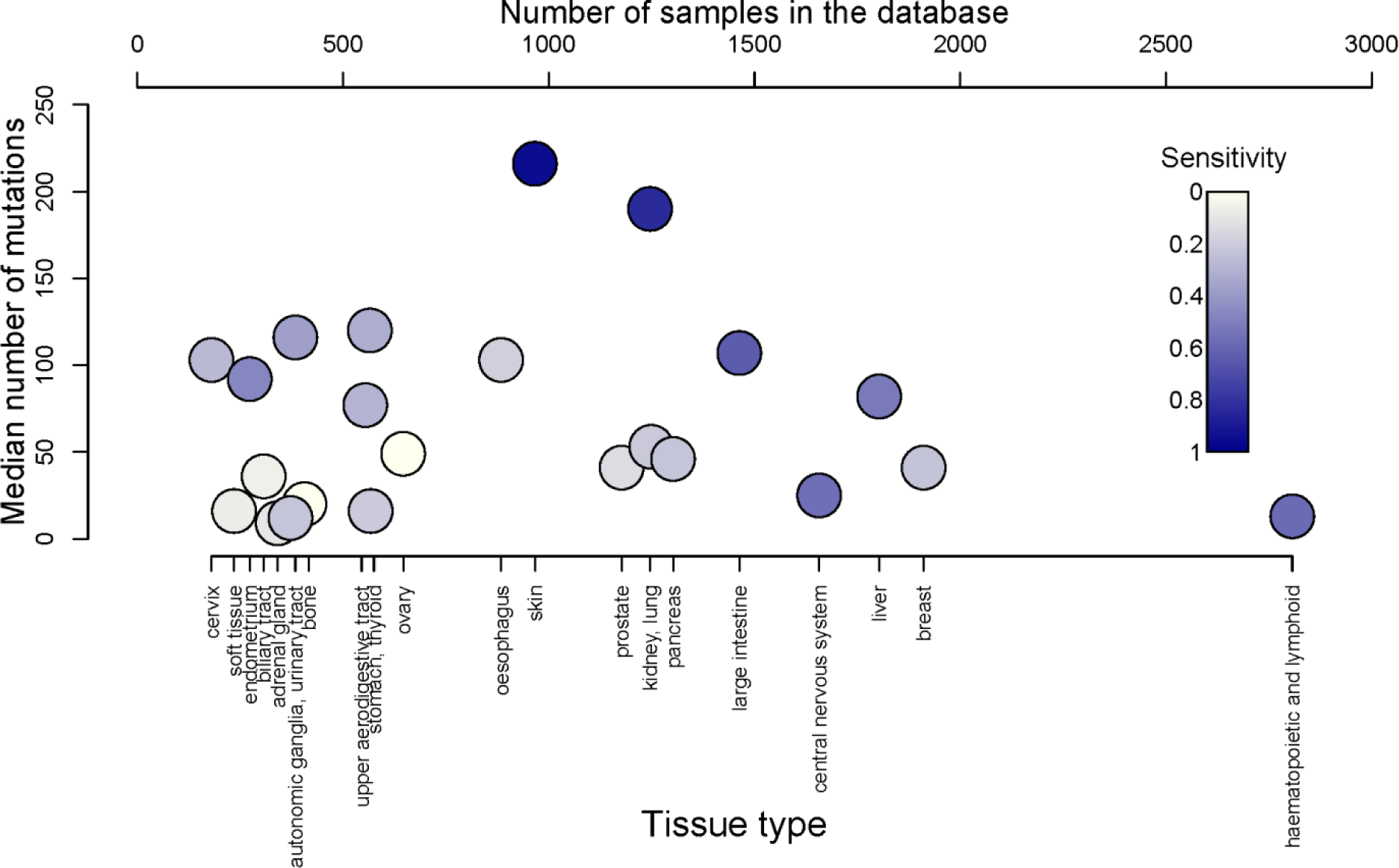
Sensitivity of detection of the 23 cancer types with their amino acid gain/loss data in the quartiles of most mutated samples with a machine learning algorithm.

### Associations of cancer types according to their amino acid gain/loss profiles

We analyzed associations of cancer types according to their amino acid gain/loss profiles (Figure 8). We found, that prostate cancer clusters most closely with pancreatic cancer, followed by cancers of central nervous system and blood. Cancer of autonomic ganglia clusters very closely with that of adrenal gland. Similarly, stomach and large intestine cancers cluster together. Unexpectedly, bone tissue cancer clusters tightly with that of biliary tract. The finding may be attributed to the small sample numbers of the cancer types in the database. The similarity of amino acid gain/loss profiles of cancers of cervix and urinary tract may be due to human papillomavirus being also active in urinary tract cancer^14^ and therefore introducing a similar nucleotide transformation signature.^2^ Breast and skin cancers are grouped in the cluster with cervical and urinary tract cancers. However, the distance between skin cancer and other cancer types in the cluster is high. Skin cancer is different from all the other tissue types in terms of high loss of proline and glycine. Furthermore, it is the only cancer type that simultaneously loses His, Met and Trp.

**Figure 8.**
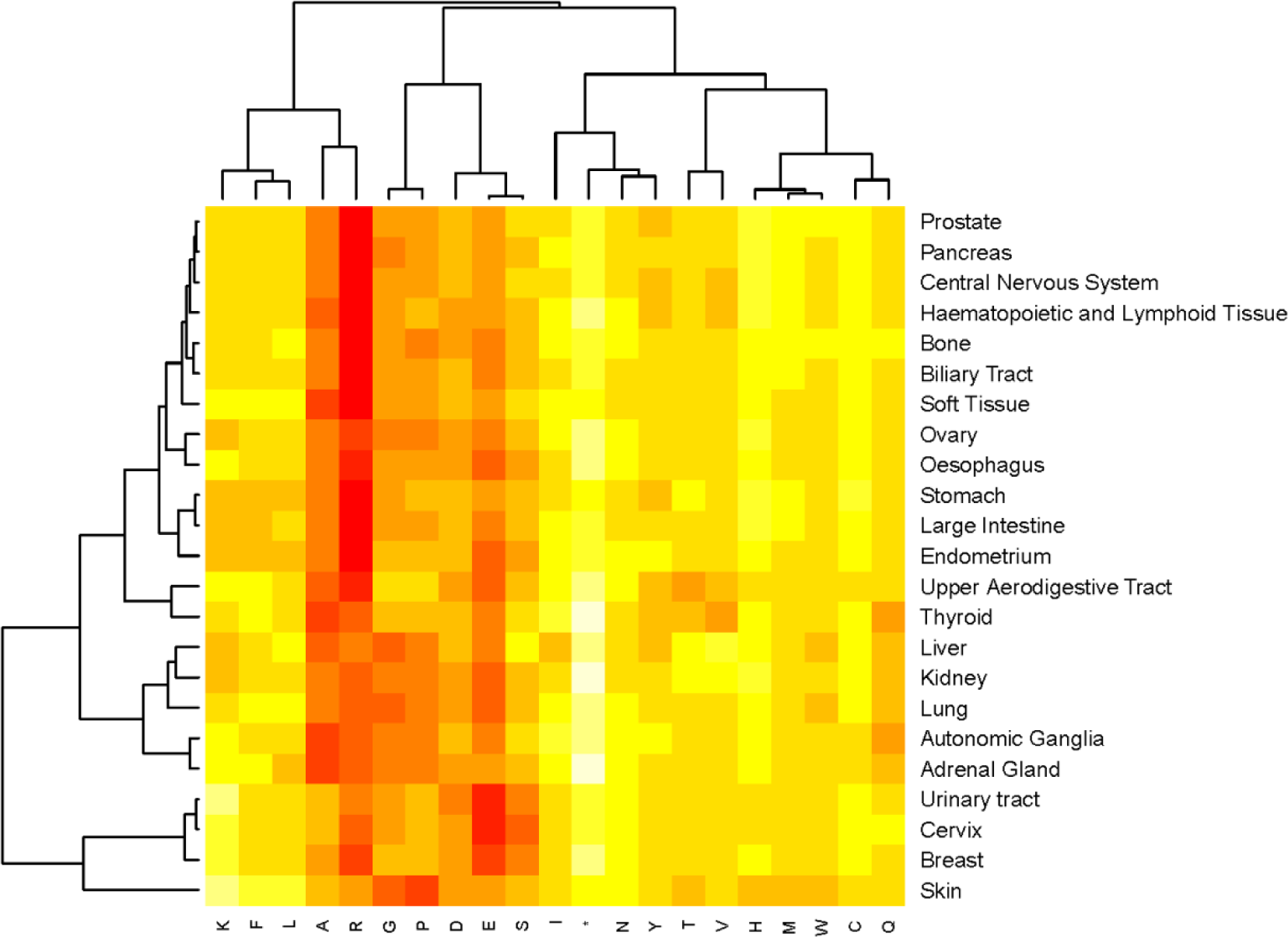
Heatmap of amino acid gain/loss profiles in 23 cancer types.

## Discussion

Cancer is associated with accumulation of mutations. During an organism’s lifetime, its DNA is constantly subjected to exogenous and endogenous damaging agents, with single nucleotide substitutions being the most common type of damage. Coding single nucleotide substitutions ultimately cause replacements of amino acids in protein sequences or, in case of transformations into stop codons, truncation events. We have shown that the net outcome of single nucleotide substitutions across multiple cancer types is a change of amino acid frequencies in the composition of proteins that makes the somatic evolution in cancer asymmetric and not in equilibrium.^4^ Six amino acids (Ala, Asp, Gln, Gly, Ser, and Arg) are irreversibly lost and the rest of amino acids are gained in a majority of cancer types. Largely, the somatic evolution in cancer shows prominent similarities with evolution of the proteome that accompanies increasing complexity of organisms.^15^ In the both processes, Pro, Ala, Gly and Glu are consistently lost, and Cys, Met, His and Phe are strongly accrued. Strikingly, while Arg did not show a universal trend either to be gained or lost in phylogenetic evolution,^15^ it is the most prominent irreversible loss in somatic evolution in cancer. We have shown that the unusual loss of Arg is associated with substitutions in the four codons out of the six coding for the amino acid that contain CG sequence in their composition. The extraordinary targeting of the CG sequence-containing codons for Arg is conceivably explained by spontaneous deamination of methylated cytosines in CpG sequences,^7^ as the CG sequence-containing codons for Ala, Pro, Thr and Ser also demonstrate moderately elevated mutation rates.^4^ DNA methylation is an epigenetic modification that increases with age^16^ and is believed to play a role in cancer.^17^ Presumably, loss of Arg is counteracted with a strong pressure of negative selection in phylogenetic evolution, while negative selection is largely absent for coding substitutions in cancer.^18^ Conversely, as evolution of species proceeds through germ cells originating from young organisms, fewer methylation sites may be present in them, and therefore fewer deamination events. However, the preponderance of irreversible loss of Arg in cancer deserves further analysis. Additionally, there are a number of other differences between somatic evolution in cancer and phylogenetic evolution. While Ser is reported to be vigorously gained in phylogenetic evolution, it is mostly lost in mutagenesis in cancer, with several tissues demonstrating very high losses of the amino acid (Figure 1). Furthermore, Lys is weakly lost in evolution of species, but predominantly gained in somatic evolution in cancer, mostly in the E>K substitution. Tyr and Trp accumulate in proteins in most cancer types, with few exceptions, but were not shown to accumulate consistently in evolution of species despite being evolutionary young amino acids.^15^

While the general tendencies of the asymmetric gains and losses of amino acids are largely consistent across 23 human cancer types in our analysis, their scopes vary significantly due to differences in mutational processes among the cancer types. Each type of either DNA damage or deficient DNA repair or replicative component has its own predilection for specific nucleotides, which can produce recognizable patterns of mutagenesis.^1,2,19^ The mutational signatures of a tumor may provide information on routes and mechanisms leading to cancer in each cancer type.^20^ We show here that at least several specific patterns of nucleotide alterations produce distinct amino acid gain/loss profiles in some cancer types, where gains and losses of some but not other amino acids cause marked changes in the proteome composition (Figure 3).

Furthermore, our analysis reveals the extent of similarity of amino acid gain/loss profiles in samples within a same cancer type. Those cancers that are predominantly associated with environmental exposures demonstrate considerably more homogeneity between samples, as seen from their higher correlation scores of amino acid gains and losses. For instance, the damage to DNA caused by ultraviolet light ensures virtually identical changes in amino acid composition in proteins of all skin cancer samples in the COSMIC database. Similarly, activities of the APOBEC family of cytidine deaminases in counteracting human papillomavirus load cause largely uniform amino acid alterations in the majority of cervical cancer samples. Likewise, mutational processes in cancers of liver and large intestine that show high correlation scores in our analysis are at least partly attributed to deleterious effects of external factors. High correlation scores in endometrial cancer may be expained by mutations in *POLE* and *POLD1* genes coding for DNA polymerases Pol δ and Pol ∊ in a few hypermutated tumor samples. The analysis suggests that the more complex assortment of mutational processes in some cancer types leads to weaker correlations of gains and losses of amino acids. The least amino acid gain/loss homogeneity was found in cancers of kidney, autonomic ganglia, ovary, bone or central nervous system. However, with mutation data on large numbers of samples it may be possible to cluster tumors within the cancer types according to their amino acid gain/loss profiles and potentially obtain information about underlying mutational processes. A machine learning algorithm using amino acid gain/loss signatures of the cancer types was able to attribute some cancer types more accurately than others, with skin cancer being the topmost correctly assigned. We suppose that a combination of the distinctivity of the amino acid gain/loss signature of a cancer type and/or its homogeneity in the available samples, the median number of mutations and the number of samples in the database influenced the ability of the algorithm to recognize cancer types.

While it takes a small number of driver mutations to cause and sustain cancer, the accompanying passenger mutations if present in high numbers may result in distinct changes of the composition of the proteome in different cancer types and therefore elevated metabolic demands for some of *de novo* gained amino acids. Furthermore, though not selected specifically for individual proteins or residue sites, some of changes may increase fitness of the cancer cell. The almost universal *de novo* introduction of Cys and His residues into the proteins may strengthen their antioxidant and metal-binding capacities. Recent data indicate that introduction of His residues in place of Arg in proteins in cancer confers an ability of pH sensing not seen with wild-type proteins.^21^ Also, the asymmetric gains and losses of chemically different amino acids may change chemical properties of the proteome. For instance, the gains of the three hydrophobic acids Phe, Leu and Ile may confer greater hydrophobicity to proteins in skin cancer, but for many cancer types actual shifts in proteome properties probably depend on the numbes of copies of affected protein molecules, as both acidic residues of Asp and Glu and basic residues of Arg are lost simultaneously. Finally, investigation of proteome alterations may contribute to the expanding realm of knowledge of underlying mutational processes in cancer.

The present analysis of amino acid gains and losses in proteins in cancer is solely based on the data on single nucleotide substitutions. The tendencies of gains and losses of amino acids may be more pronounced when affected proteins are present in high numbers in the cancer cell, or less prominent when mutation-carrying proteins have insufficient stability or are low expressed. Experimental analyses are needed to evaluate the actual effect of the mutation-induced gains and losses of amino acids on the composition of cellular proteins in cancer. Similarly, as many mutational processes are also operative in noncancerous tissue, it would be informative to determine to what extent the amino acid gain/loss tendencies effect composition of proteins in normal tissue.

### Supporting Information

Figures S1−S11 (PDF)

Table S1-S2 (XLSX)

AUTHOR INFORMATION

### Funding Sources

The author has no support or funding to report.

